# Disrupted Decision-Making: EcoHIV Inoculation Following a History of Cocaine Use

**DOI:** 10.1101/2022.05.13.491868

**Authors:** Kristen A. McLaurin, Hailong Li, Charles F. Mactutus, Steven B. Harrod, Rosemarie M. Booze

## Abstract

Independently, chronic cocaine use and HIV-1 viral protein exposure induce neuroadaptations in the frontal-striatal circuit; how the frontal-striatal circuit responds to HIV-1 infection following chronic drug use, however, has remained elusive. After establishing a history of both sucrose and cocaine self-administration, a pretest-posttest experimental design was utilized to evaluate preference judgment, a simple form of decision-making dependent upon the integrity of frontal-striatal circuit function. During the pretest assessment, male rats exhibited a clear preference for cocaine, whereas female animals preferred sucrose. Two posttest evaluations (3 Days and 6 Weeks Post Inoculation) revealed that, independent of biological sex, inoculation with chimeric HIV (EcoHIV), but not saline, disrupted decision-making. Prominent structural alterations in frontal-striatal circuit dysfunction were evidenced by synaptodendritic alterations in pyramidal neurons in the medial prefrontal cortex. Thus, the EcoHIV rat affords a biological system to model how the frontal-striatal circuit responds to HIV-1 infection following chronic drug use.

## INTRODUCTION

In 2020, approximately 5.2 million individuals living in the United States reported having used cocaine in the past year, whereby nearly 1.3 million of these individuals met the *Diagnostic and Statistical Manual of Mental Disorders* (5^th^ ed.; DSM-5) criteria for cocaine use disorder (SAMHSA, 2020). Intravenous (IV) drug use significantly increases risky behaviors (e.g., needle sharing, unprotected sex, trading sex for drugs or money) associated with contracting infectious diseases, including human immunodeficiency virus type 1 (HIV-1; Hudgins et al., 1995). Indeed, individuals who inject drugs have a significantly greater (i.e., up to 35 times) risk of acquiring HIV-1 (Chaisson et al., 1989; McCoy et al., 2004; UNAIDS, 2022). Independently, chronic cocaine use or HIV-1 viral protein exposure induce significant neurochemical and structural adaptations in the frontal-striatal circuit; how the frontal-striatal circuit responds to HIV-1 infection following a history of cocaine use, however, has remained elusive (for review, Illenberger et al., 2020).

Originally proposed by Alexander et al. (Alexander et al., 1986; Alexander and Crutcher, 1990; Alexander, 1994), the frontal-striatal circuits include five parallel segregated circuits connecting selected cortical areas to the basal ganglia and thalamus. Of particular relevance to substance use disorders is the frontal-striatal circuit linking the prefrontal cortex (PFC), nucleus accumbens (NAc), and ventral tegmental area (VTA). Specifically, excitatory glutamatergic afferents from the PFC directly target the NAc (Haber et al., 1995) and VTA (Sesack et al., 1989; Sesack and Pickel, 1992). Dopaminergic projections from the VTA innervate both the PFC (Oades and Halliday, 1987) and NAc (Swanson, 1982). Inhibitory gamma-aminobutyric acid (GABA) afferents from the NAc innervate the PFC via the mediodorsal nucleus of the thalamus (Ray and Price, 1993) and target dopaminergic neurons in the VTA (Watabe-Uchida et al., 2012).

Chronic use of the psychostimulant cocaine induces neurochemical and structural neuroadaptations in the frontal-striatal circuit. Motivationally relevant events, including acute cocaine use, induce activation of dopaminergic neurons in the VTA leading to a rapid potentiation of dopamine (DA) in the NAc (Bradberry et al., 1993; Pontieri et al., 1995; Yuen et al., 2021). Across repeated exposures to a motivationally relevant event (i.e., as would be observed with a history of cocaine use), however, dopamine function decreases in both the NAc (e.g., Volkow et al., 1990; Volkow et al., 1997; Ferris et al., 2012; Willuhn et al., 2014) and PFC (Briand et al., 2008); alterations in glutamate homeostasis also develop (for review, Kalivas, 2009). Structural alterations to neurons and their associated dendritic spines, which occur following a history of cocaine self-administration (Robinson et al., 2001), may reflect the reorganization of synaptic connectivity within the frontal-striatal circuit.

In a similar manner, chronic HIV-1 viral protein exposure reorganizes the frontal-striatal circuit resulting in the establishment of neurocognitive impairments and affective alterations. Specifically, chronic HIV-1 viral protein exposure induces a hypodopaminergic state (for review, McLaurin et al., 2021a; e.g., Kumar et al., 2009; Denton et al., 2019) and glutamatergic dysregulation (for review, Vázquez-Santiago et al., 2014), as well as prominent synaptodendritic neuronal injury (e.g., Moore et al., 2006; Roscoe et al., 2014; McLaurin et al., 2019; Festa et al., 2020; Speidell et al., 2020). Fundamentally, neurochemical and structural alterations in the frontal-striatal circuit have been associated with neurocognitive (Chang et al., 2008; Kumar et al., 2011) and affective (Paul et al., 2005; Kamat et al., 2014) alterations in HIV-1 seropositive individuals. Given the prominent frontal-striatal dysfunction resulting from either cocaine dependence or HIV-1, there remains a fundamental need to develop a biological system to model how the frontal-striatal circuit responds to HIV-1 infection following a history of drug dependence.

In 2005, Potash et al. reported the development of a novel chimeric HIV-1 virus construct (EcoHIV) to model active HIV-1 infection in mice. The EcoHIV construct preserves all of the cis-regulatory elements of HIV-1, but replaces the coding region of gp120 with gp80 from ecotropic murine leukemia virus (MLV; Potash et al., 2005). Host’s cells are infected with ecotropic MLV via the cationic amino acid transporter (CAT-1; Kim et al., 1991, Wang et al., 1991). Critically, EcoHIV infection recapitulates many of the clinical features of HIV-1, including lymphocyte, macrophage, and microglia infection, neurocognitive impairments, and synaptodendritic alterations (e.g., Geraghty et al., 2017; Gu et al., 2018; Kelschenbach et al., 2019; Li et al., 2021a). The recent extension of the EcoHIV model of HIV-1 infection to rats (Li et al., 2021a; 2021b), which are more commonly used in studies of substance use disorders and neurocognitive impairments, may afford a biological system to investigate comorbid cocaine use and HIV-1.

Thus, in the present study, after establishing a drug dependent phenotype, male and female rats were inoculated with either EcoHIV or saline. A pretest-posttest experimental design (Figure 1) was utilized to evaluate preference judgment, a simple form of decision-making (Fellows, 2007), using a concurrent choice self-administration experimental paradigm. Anatomically, pyramidal neurons, and their associated dendritic spines, in the medial PFC (mPFC) were evaluated as an index of frontal-striatal circuit function. Prominent disruptions in decision-making and structural reorganization of the frontal-striatal circuit in the EcoHIV rat affords an instrumental model system to investigate how the frontal-striatal circuit responds to HIV-1 infection following a history of cocaine use.

**Figure 1.**
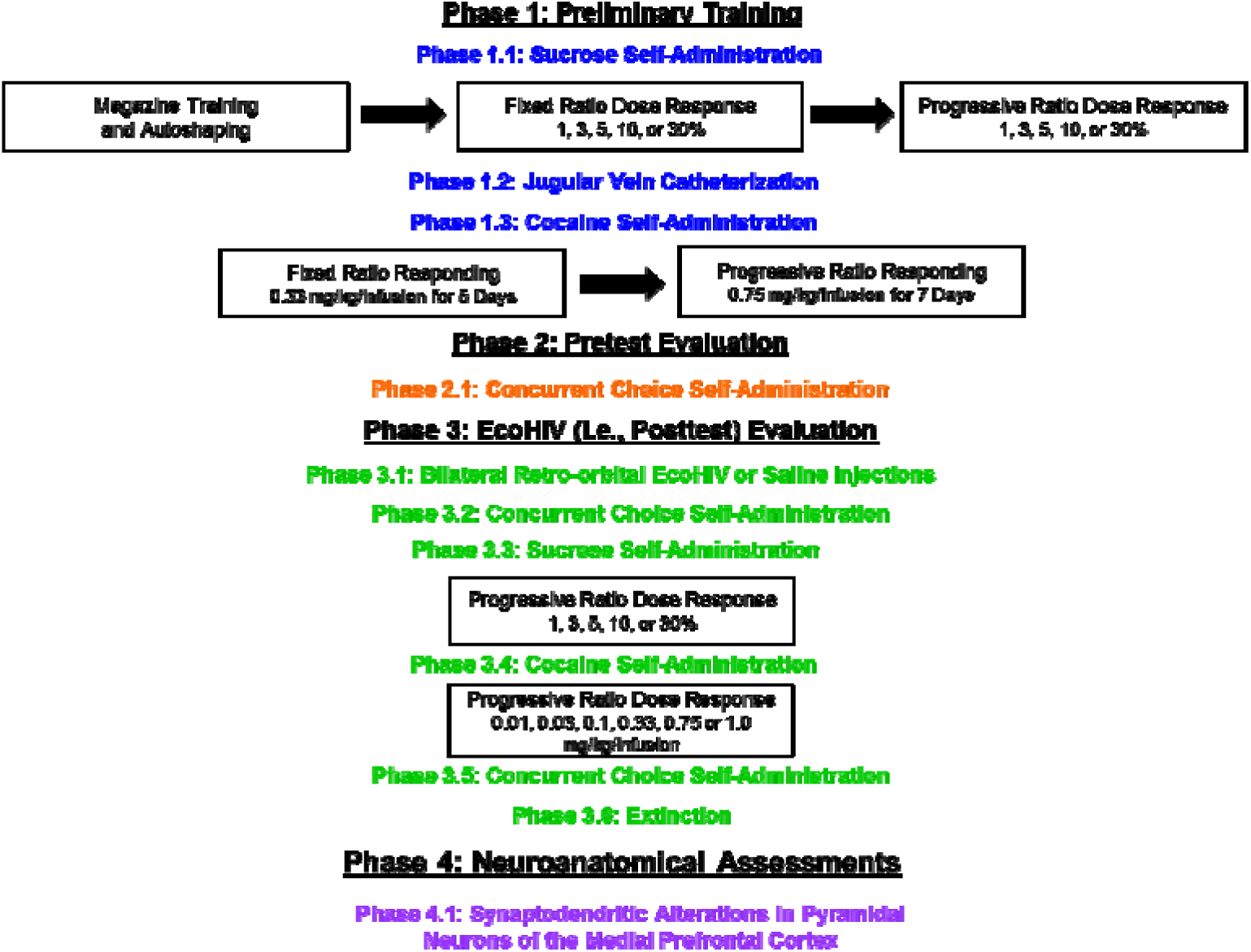
Pretest-Posttest Experimental Design Schematic.

## RESULTS

### Phase 1.3: Escalation of cocaine-maintained responding supports the development of a drug dependent phenotype

Drug dependence is characterized, at least in part, by escalation of drug self-administration (for review, Edwards and Koob, 2013). Male and female animals were trained to self-administer cocaine using both fixed ratio (FR; 5 Days, 0.33 mg/kg/infusion) and progressive ratio (PR; 7 Days, 0.75 mg/kg/infusion) schedules of reinforcement. Animals, independent of biological sex, exhibited a profound escalation of cocaine self-administration under a PR schedule of reinforcement. Specifically, across the seven PR test sessions, animals exhibited a linear increase in the number of active lever presses (Supplementary Figure 1A; Main Effect: Day, *F*(6, 216)=3.7, *p*≤0.01; Best Fit: First Order Polynomial, *R*^2^≥0.85) and the number of cocaine reinforcers (Supplementary Figure 1B; Main Effect: Day, *F*(6, 216)=8.1, *p*≤0.01; Best Fit: First Order Polynomial, *R*^2^≥0.94) supporting the development of a drug dependent phenotype.

### Phase 2.1: Prominent sex differences in preference judgment were observed during the pretest evaluation

Following a history of both sucrose and cocaine self-administration, male and female animals were evaluated in a concurrent choice self-administration experimental paradigm (5 Days) to evaluate preference judgment. Animals responded on a FR-1 schedule of reinforcement for either 5% sucrose (weight/volume (w/v)) or 0.33 mg/kg/infusion cocaine.

Prominent sex differences in preference judgment, a simple form of decision-making, were observed during the pretest evaluation (Figure 2; Sex x Reinforcer x Day Interaction, *F*(4, 104)=4.8, *p*≤0.01). Male animals established a clear preference for cocaine (Best Fit: Horizontal Line), relative to sucrose (Best Fit: Segmental Linear Regression, *R*^2^ ≥ 0.94); a preference judgment that was maintained across the five day testing period (Main Effect: Reinforcer, *F*(1,17)=5.7, *p*≤0.03). In sharp contrast, female animals developed a preference for sucrose (Best Fit: Segmental Linear Regression, *R*^2^≥0.99), relative to cocaine (Best Fit: Horizontal Line), across the testing period (Day x Reinforcer Interaction, *F*(4,46)=4.7, *p*≤0.01).

**Figure 2.**
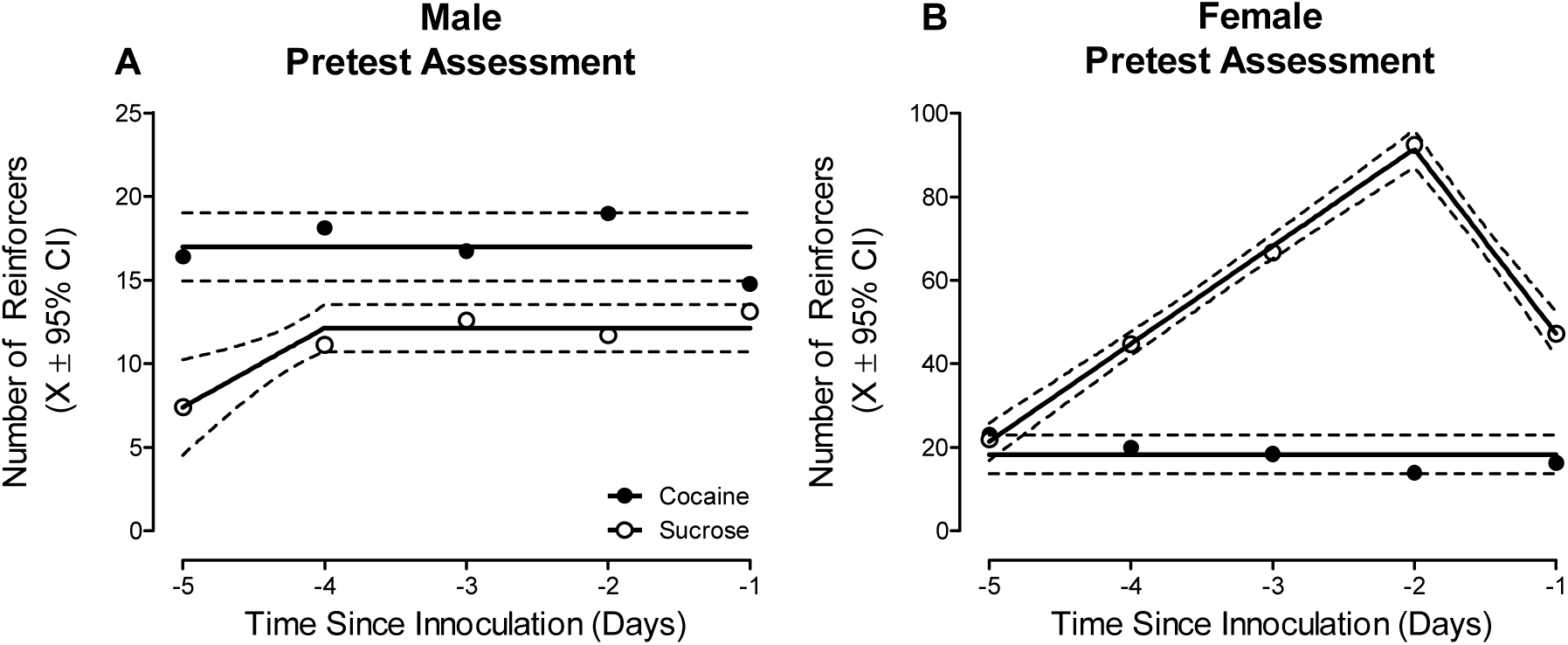
A concurrent choice self-administration paradigm was utilized to evaluate preference judgment as a simple form of decision-making. During the pretest assessment, male (**A**) animals consumed a greater number of cocaine, relative to sucrose, reinforcers; observations which indicate a clear, consistent preference for cocaine. Female (**B**) animals, however, developed a preference for sucrose, relative to cocaine, across the five day testing period.

### Phase 3.1: Bilateral retro-orbital inoculation of EcoHIV induced significant HIV infection in the medial prefrontal cortex; infection which is primarily harbored in microglia

Animals were assigned to receive bilateral retro-orbital injections of either EcoHIV or saline based on measurements from preliminary training (Phase 1). After the completion of experimentation, immunohistochemical staining was utilized for two complementary purposes, including 1) to quantify the amount of EcoHIV infection present in the mPFC; and 2) to establish the predominant cell type harboring EcoHIV infection.

All animals, independent of biological sex, exhibited significant (Range: 699-2958) EcoHIV-EGFP expression in the mPFC at sacrifice (i.e., approximately eight weeks post inoculation; Figure 3). No statistically significant sex differences in the number of EcoHIV-EGFP signals were observed (*p*>0.05). Second, EcoHIV-EGFP and Iba1, a microglial marker, were highly co-localized in the mPFC supporting microglia as the predominant cell type for EcoHIV expression; observations which are consistent with previous reports in HIV-1 seropositive individuals (Ko et al., 2016) and the EcoHIV rat (Li et al., 2021a, 2021b).

**Figure 3.**
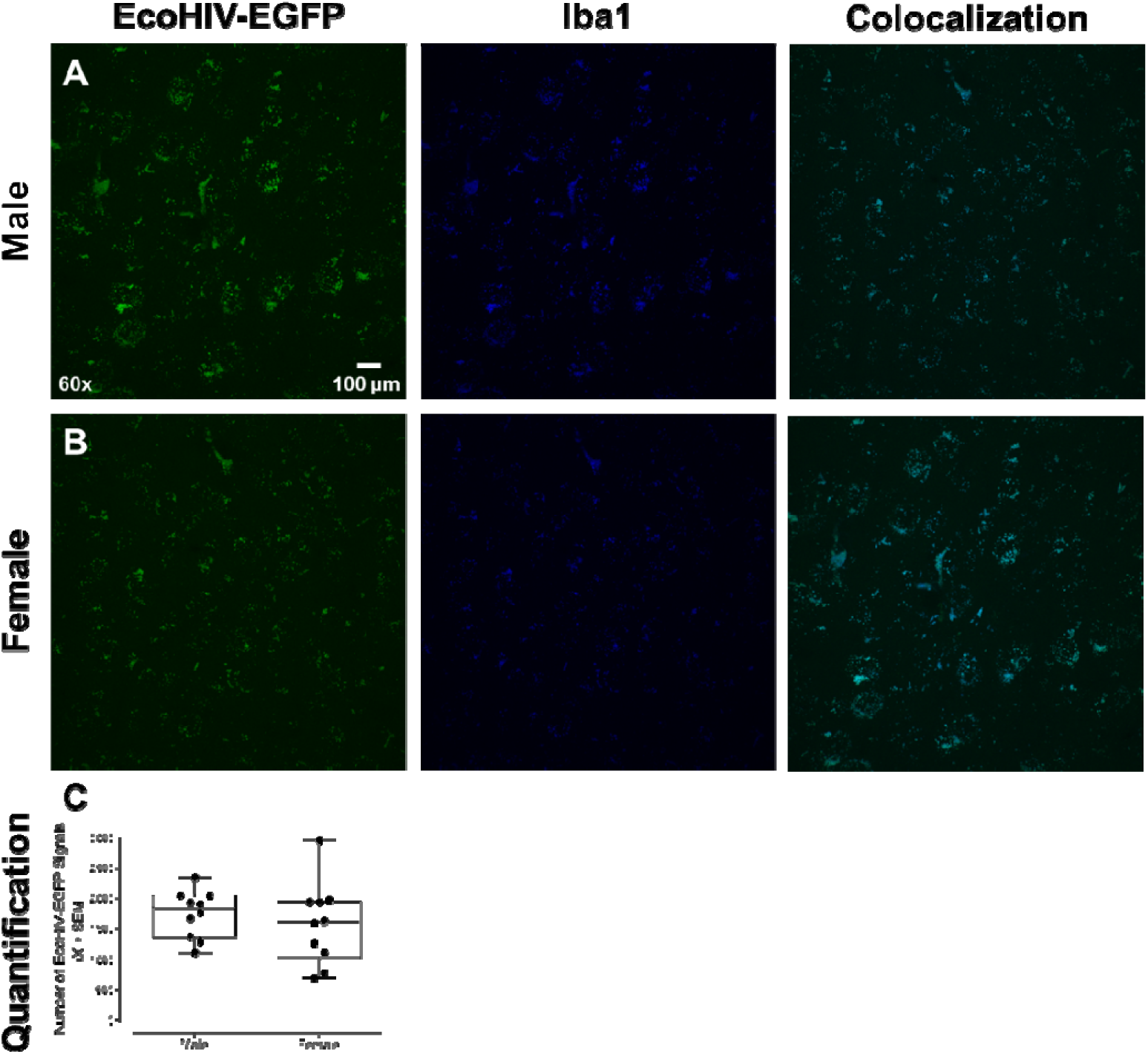
Representative images of EcoHIV-EGFP, Iba1, a marker of microglia, and their co-localization in the medial prefrontal cortex (mPFC) in male (**A**) and female (**B**) animals. The number of EcoHIV-EGFP signals were quantified; no statistically significant sex differences in the number of EcoHIV-EGFP signals were observed (**C**).

### Phases 3.2 to 3.5: EcoHIV inoculation disrupted preference judgment, but not reinforcing efficacy, during posttest evaluations

Beginning three days after inoculation, EcoHIV and saline animals were reevaluated in the concurrent choice self-administration experimental paradigm for five days. Inoculation with EcoHIV disrupted preference judgment in both male and female animals (Figure 4; Treatment x Sex x Reinforcer Interaction, *F*(1, 31)=7.2, *p*≤0.01). Specifically, male EcoHIV animals failed to exhibit a preference for either sucrose or cocaine (Best Fit: Global Horizontal Line; Test of Difference in Means: *p*>0.05). Female EcoHIV animals exhibited an initial preference for sucrose (Best Fit: Segmental Linear Regression, *R*^2^≥0.97), relative to cocaine (Best Fit: Segmental Linear Regression, *R*^2^≥0.96); a preference which dissipated during the first posttest evaluation. In sharp contrast, saline male and female rats maintained their clear preference for cocaine (Best Fit: Horizontal Line; Test of Difference in Means, *F*(1,8)=52.4, *p*≤0.01) or sucrose (Best Fit: Horizontal Line; Test of Difference in Means, *F*(1,8)=187.9, *p*≤0.01), respectively.

**Figure 4.**
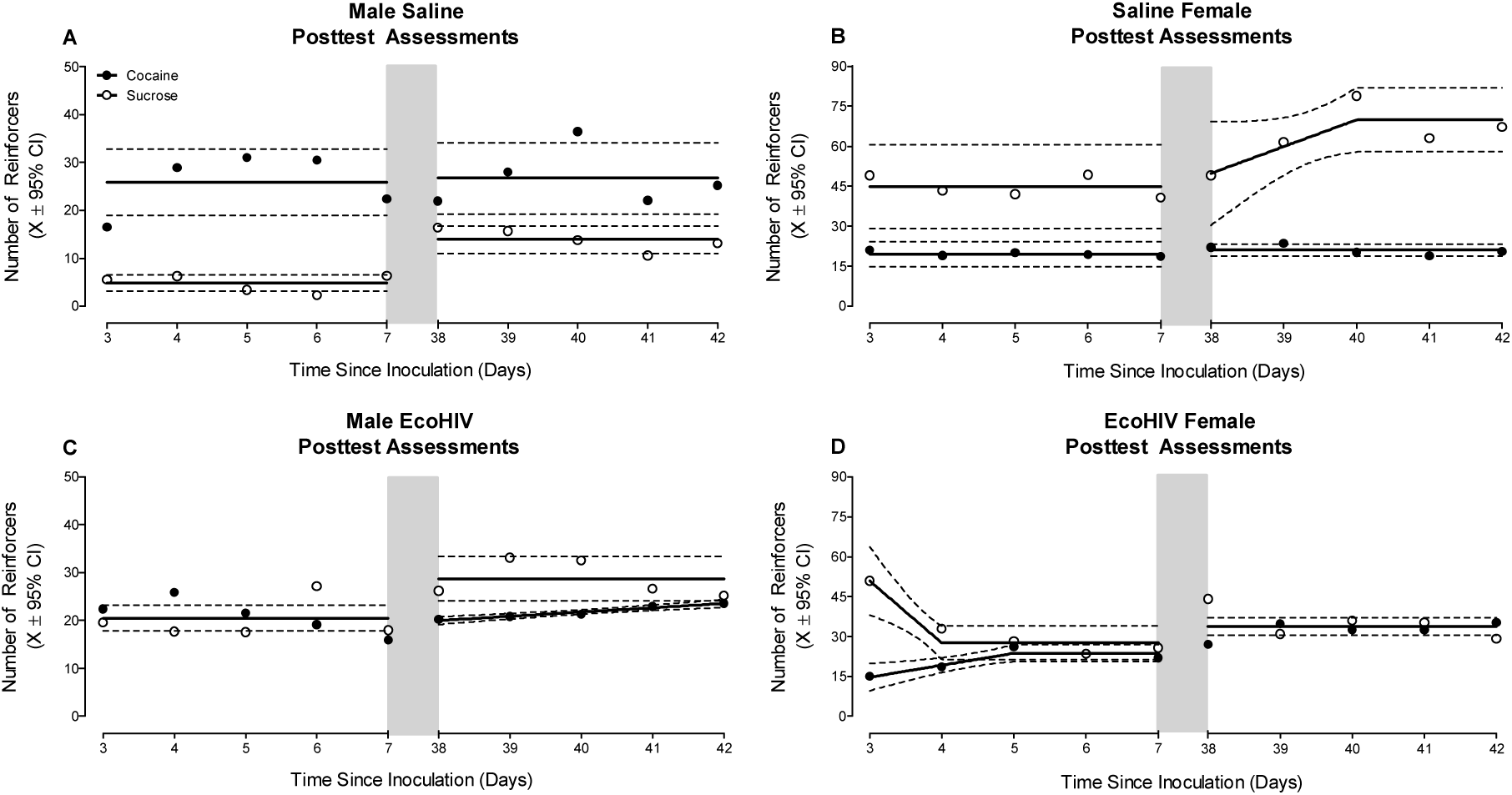
Beginning three days after EcoHIV or saline inoculation, animals were reevaluated in the concurrent choice self-administration experimental paradigm. A second posttest evaluation was conducted approximately six weeks after inoculation. The light gray bar indicates the time period between the first and second posttest evaluations. Saline male (**A**) and female (**B**) rats maintained their clear preference for cocaine or sucrose, respectively, during both posttest evaluations; a preference that was consistent with observations during the pretest assessment. In sharp contrast, EcoHIV inoculation disrupted preference judgment in both male (**C**) and female (**D**) animals. Specifically, the disruption of preference judgment in EcoHIV animals was characterized by either the failure to exhibit a clear preference for either reinforcer or a change in preference judgment across time.

A second posttest evaluation of preference judgment was conducted approximately six weeks after EcoHIV or saline inoculation. Disruption of preference judgment in both male and female EcoHIV animals persisted (Figure 4; Treatment x Sex x Reinforcer Interaction, *F*(1,32)=32.0, *p*≤0.01). Specifically, male EcoHIV animals preferred sucrose (Best Fit: Horizontal Line) relative to cocaine (Best Fit: First-Order Polynomial, *R*^2^≥0.95); a sharp contrast to the maintained preference for cocaine observed in male saline rats (Best Fit: Horizontal Line; Test of Difference in Means, *F*(1,78)=8.0, *p*≤0.01). Female EcoHIV rats failed to exhibit a preference for either sucrose or cocaine (Best Fit: Horizontal Line; Test of Difference in Means, *p*>0.05), whereas female saline rats again displayed a clear preference for sucrose (Best Fit: Segmental Linear Regression, *R*^2^≥0.85) over cocaine (Best Fit: Horizontal Line).

Notably, the disruption of preference judgment observed following EcoHIV inoculation occurred in the absence of any changes in the reinforcing efficacy of either sucrose or cocaine (Supplementary Figure 2). Specifically, sucrose concentration (1, 3, 5, 10, or 30% w/v; Phase 3.3) and cocaine dose (0.01, 0.03, 0.1, 0.33, 0.75, or 1 mg/kg/infusion; Phase 3.4) were systematically manipulated under a PR schedule of reinforcement to evaluate reinforcing efficacy (Richardson and Roberts, 1996). EcoHIV inoculation failed to alter the reinforcing efficacy of either sucrose (*p*>0.05; Best Fit: Global First-Order Polynomial, *R*^2^≥0.83 and *R*^2^≥0.79 for the Number of Active Lever Presses and Sucrose Reinforcers, respectively) or cocaine (*p*>0.05; Best Fit: Global First-Order Polynomial, *R*^2^≥0.94 and *R*^2^≥0.91 for the Number of Active Lever Presses and Cocaine Reinforcers, respectively) during the PR dose response assessments. Thus, infection with EcoHIV induced a prominent disruption in preference judgment, but not reinforcing efficacy.

### Phase 3.6: EcoHIV animals exhibited blunted extinction learning following removal of the preferred reinforcer

Following completion of the second posttest evaluation, each animals’ preferred reinforcer (i.e., Cocaine or Sucrose) was individually determined and replaced (i.e., Cocaine with Saline; Sucrose with Water) to evaluate extinction learning in the concurrent choice self-administration experimental paradigm.

EcoHIV inoculation blunted extinction learning relative to saline inoculation; the pattern of which was dependent upon the reinforcer removed (Treatment x Day x Reinforcer Removed Interaction with a Prominent Quadratic Component, *F*(1,30)=6.6, *p*≤0.02). Following the removal of cocaine, EcoHIV animals failed to extinguish to the same level as saline animals (Figure 5A; Main Effect: Treatment, F(1,22)=6.4, *p*≤0.02). Observations of blunted cocaine extinction learning in EcoHIV rats were confirmed using complementary regression analyses (Best-Fit Function: One-Phase Decay, *R*^2^s≥0.92; Differences in the Parameters of the Function *F*(3,10)=10.6, *p*≤0.01). The removal of sucrose also induced a blunted rate of extinction in EcoHIV, relative to saline, animals (Figure 5B; Day x Treatment Interaction with a Prominent Quadratic Component, *F*(1,8)=6.6, *p*≤0.04). Although sucrose extinction learning was well-described by a one-phase decay in both EcoHIV (*R*^2^s≥0.97) and saline (*R*^2^s≥0.97) animals, significant differences in the parameters of the function were observed (*F*(3,10)=11.9, *p*≤0.01). Taken together, removal of the preferred reinforcer revealed the generality of PFC deficits in the EcoHIV rat by extending neurocognitive impairments to include blunted extinction learning.

**Figure 5.**
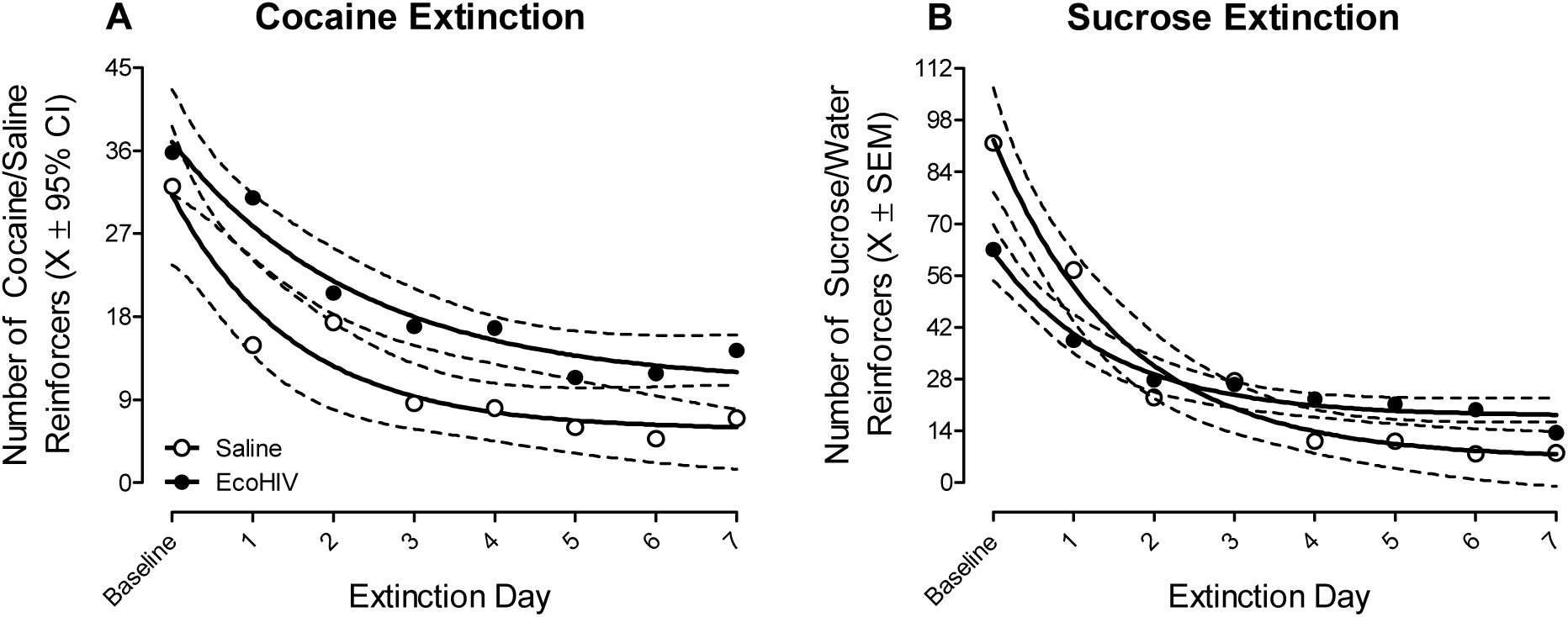
Following completion of the second posttest evaluation, each animals’ preferred reinforcer (i.e., Cocaine or Sucrose) was individually determined and replaced (i.e., Cocaine with Saline; Sucrose with Water) to evaluate extinction learning in the concurrent choice self-administration experimental paradigm. Blunted extinction learning was observed after EcoHIV, but not saline, inoculation. Specifically, following the removal of either cocaine (**A**) or sucrose (**B**), EcoHIV rats failed to decrease their responding to the same levels as saline animals.

### Phase 4: Inoculation with EcoHIV induced prominent synaptodendritic alterations in pyramidal neurons, and their associated dendritic spines, in the medial prefrontal cortex

Following the completion of experimentation, pyramidal neurons in the mPFC, and their associated dendritic spines, were labeled using a ballistic labeling technique (Li et al., 2020).

Morphological characteristics of pyramidal neurons, and their associated dendritic spines, were examined along the apical dendrite using sophisticated neuronal reconstruction software.

A centrifugal branch ordering scheme was utilized to examine dendritic branching complexity and synaptic connectivity along the apical dendrite from Branch Orders 1 to 12. Infection with EcoHIV failed to alter dendritic branching complexity in pyramidal neurons in the mPFC (Figure 6A; *p*>0.05), evidenced by a global Gaussian fit (*R*^2^≥0.97) for the number of dendritic branches at each branch order. With regards to synaptic connectivity, EcoHIV inoculation induced a profound population shift in the location of dendritic spines along the apical dendrite (Figure 6B; Genotype x Branch Order Interaction, *F*(11,362)=47.1, *p*≤0.01). Specifically, EcoHIV, relative to saline, animals exhibited an increased relative frequency of dendritic spines on distal branches along the apical dendrite. The factor of biological sex altered the magnitude, but not the pattern, of the population shift in the location of dendritic spines along the apical dendrite (Genotype x Sex x Branch Order Interaction, *F*(11, 362)=127.8, *p*≤0.01).

**Figure 6.**
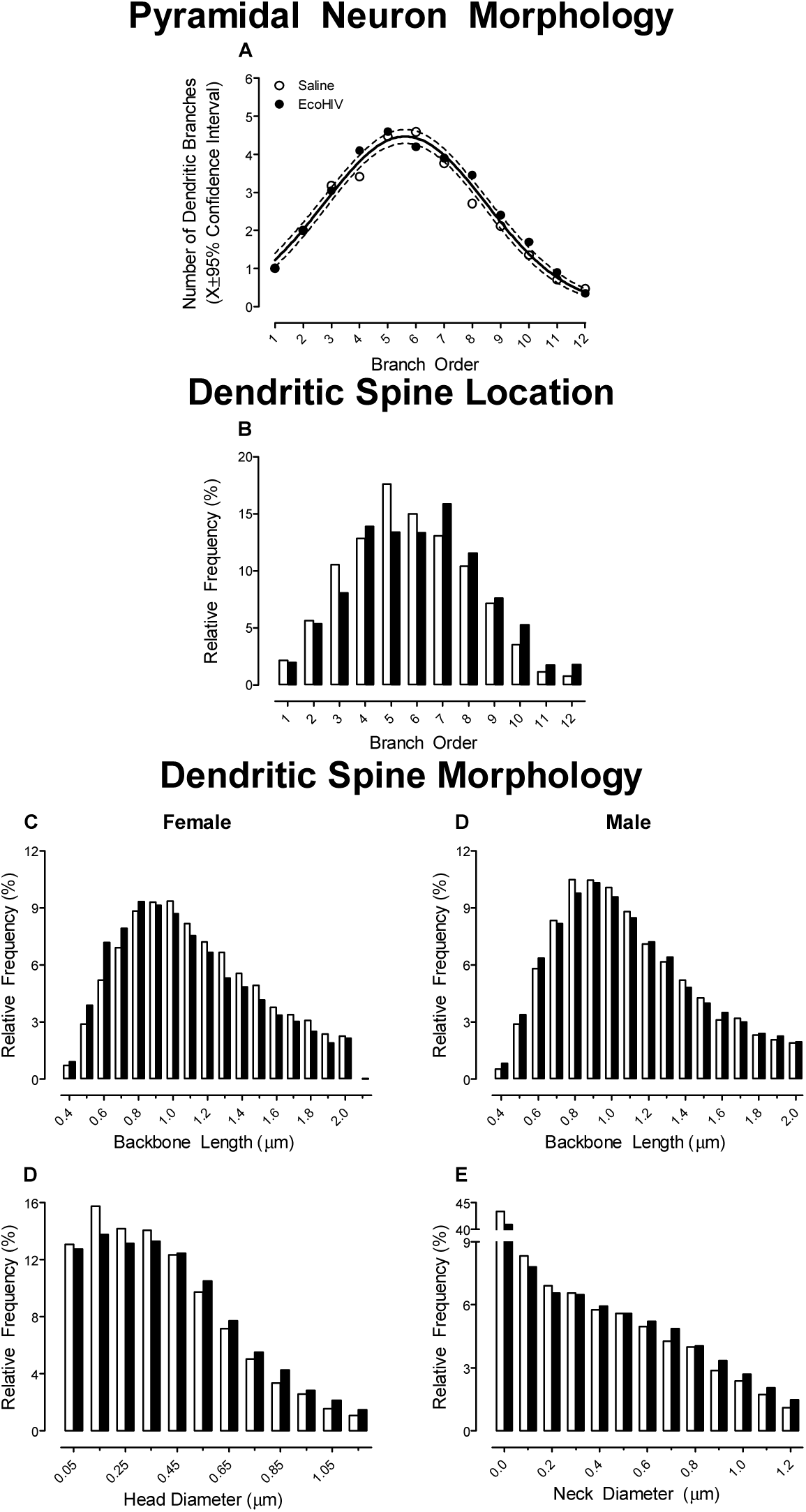
Prominent synaptic dysfunction in pyramidal neurons from layers II-III of the medial prefrontal cortex was observed following EcoHIV inoculation. (**A**) Utilization of a centrifugal branch ordering scheme to evaluate dendritic branching complexity revealed no statistically significant differences between EcoHIV and saline animals. (**B**) A profound alteration in synaptic connectivity was evidenced by a population shift towards a greater relative frequency of dendritic spines on more distal branches in EcoHIV, relative to saline, rats. (**C-E**) Morphological parameters of dendritic spines in EcoHIV animals were shifted towards a more immature phenotype, characterized by a sex-dependent alterations in backbone length and a population shift towards increased dendritic spine head and neck diameter.

In addition, three parameters, including backbone length (μm), head diameter (μm), and neck diameter were used to characterize dendritic spine morphology in EcoHIV and saline animals. A population shift in dendritic spine backbone length was dependent upon EcoHIV inoculation and biological sex (Figures 6C-D; Genotype x Sex x Bin Interaction, *F*(16, 528)=2.8, *p*≤0.01). Specifically, female EcoHIV animals displayed a prominent population shift towards shorter dendritic spines relative to female saline animals (Genotype x Bin Interaction, *F*(16, 256)=6.2, *p*≤0.01); no statistically significant genotypic alterations were observed in male animals (*p*>0.05). Furthermore, EcoHIV animals displayed a profound population shift towards increased dendritic spine head diameter (Figure 6E; Genotype x Bin Interaction, *F*(11, 363)=8.6, *p*≤0.01) and neck diameter (Figure 6F; Genotype x Bin Interaction, *F*(12, 396)=6.1, *p*≤0.01); the magnitude of which was dependent upon biological sex (Head Diameter: Genotype x Sex x Bin Interaction, *F*(11, 363)=3.7, *p*≤0.01; Neck Diameter: Genotype x Sex x Bin Interaction, *F*(12, 396)=10.4, *p*≤0.01). Collectively, EcoHIV induced a structural reorganization of the frontal-striatal circuit, evidenced by prominent alterations in synaptic connectivity and a population shift towards an immature dendritic spine phenotype in the mPFC relative to saline animals.

## DISCUSSION

EcoHIV inoculation following a history of cocaine dependence induced profound functional and structural neuroadaptations in the frontal-striatal circuit. Using a pretest-posttest experimental design, preference judgment, a simple form of decision-making, was evaluated using a concurrent choice self-administration experimental paradigm. During the pretest assessment, male rats exhibited a significantly greater number of responses (i.e., preference) for cocaine, whereas female animals preferred sucrose. Bilateral retro-orbital inoculation of EcoHIV, but not saline, disrupted decision-making and blunted extinction learning in both male and female rats. Notably, the prominent EcoHIV induced alterations in decision-making occurred in the absence of any changes in the reinforcing efficacy of either sucrose or cocaine. EcoHIV rats, independent of biological sex, also exhibited prominent structural neuroadaptations in pyramidal neurons of the mPFC evidenced by decreased synaptic connectivity and a population shift towards an immature dendritic spine phenotype. Collectively, the EcoHIV rat affords a biological system to model functional and structural frontal-striatal circuit responses to HIV-1 infection following chronic drug use.

Drug dependence is characterized, at least in part, by the transition from recreational to uncontrollable (i.e., escalation of drug use frequency and/or intensity and compulsivity) drug use (Edwards and Koob, 2013). Preclinical self-administration studies have provided a multitude of experimental paradigms to model key aspects of the drug dependence process. Specifically, rats will escalate their cocaine use frequency and/or intensity under extended access conditions (i.e., access to drug for at least six hours; Ahmed and Koob, 1998; 1999) and/or PR schedules of reinforcement (Morgan et al., 2006; McLaurin et al., 2021b, Present Study). The escalation of cocaine self-administration in preclinical biological systems also induces prominent frontal-striatal circuit dysfunction evidenced by alterations in dopamine neurotransmission (Ahmed and Koob, 2003; Willuhn et al., 2014; Bertrand et al., 2018), glutamatergic receptors (Hao et al., 2010), and glutamatergic signaling genes (Ben-Shahar et al., 2009; Ploense et al., 2017). Thus, the pronounced escalation of cocaine-maintained responding promoted in the present study supports the development of a drug dependent phenotype and dysfunction in the frontal-striatal circuit.

Animals were inoculated with EcoHIV or saline via retro-orbital injections after the development of a drug dependent phenotype to investigate preference judgment, a simple form of decision-making (Fellows, 2007). By definition, decision-making relies upon both external (e.g., position of the operant lever during forced-choice trials) and internal (e.g., reinforcer history) information to develop a motor plan. Preference judgment, more specifically, requires making a choice (i.e., decision) by comparing the relative value of options (e.g., sucrose vs. cocaine). EcoHIV inoculation induced an early and persistent disruption in preference judgment, characterized by inconsistent choices, altered preferences, and/or blunted extinction learning, relative to saline animals; findings which are consistent with those reported in the HIV-1 transgenic rat (Bertrand et al., 2018). Impairments in more complex decision-making tasks, tapped using the Iowa Gambling Task, have also been reported in HIV-1 seropositive individuals with (Martin et al., 2004) and without (Hardy et al., 2006; Iudicello et al., 2013) drug dependency. Notably, disruption in preference judgment may precede and/or underlie impairments in more complex decision-making necessitating further investigation of these fundamental neurocognitive functions.

Anatomically, decision-making relies, at least in part, on the integrity of the frontal-striatal circuit. Specifically, lesions, inactivation, or receptor knockdowns in the PFC (Bechara et al., 1998; Walton et al., 2002), NAc (Cardinal et al., 2001; Hauber and Sommer, 2009; Floresco et al., 2018), or VTA (Bernosky-Smith et al., 2018) profoundly alter complex (i.e., effort-based, risk-based) decision-making. With specific regards for preference judgment, lesions of the PFC induce prominent alterations in preference judgment, characterized by inconsistent choices (e.g., Baylis and Gaffan, 1991; Fellows and Farah, 2007; Henri-Bhargava et al., 2012); findings which resemble observations in the EcoHIV rat. More critically, altering the connections between brain regions of the frontal-striatal circuit, either via chemogenetic approaches or asymmetrical excitotoxic lesions, also disrupts complex decision-making (PFC-NAc: Hauber and Sommer, 2009) and preference judgment (PFC-NAc: Oyama et al., 2021).

Given the fundamental role of the PFC in preference judgment and its reciprocal connections with both the NAc and VTA, pyramidal neurons, and their associated dendritic spines, in layer II-III of the mPFC were examined as an index of frontal-striatal circuit function. The dendritic tree of pyramidal neurons in layers II-III of the mPFC is characterized by a single apical dendrite arising from the apex of the soma and short basilar dendrites descending from the base of the soma (for review, Spruston, 2008). An apical tuft, formed by the bifurcation of the apical dendrite, is covered with dendritic spines that serve as the primary target for synaptic inputs (Hersch and White, 1981). Examination of dendritic spines, therefore, affords a key opportunity to infer the integrity of frontal-striatal circuit function.

Prominent synaptodendritic alterations were observed in pyramidal neurons, and their associated dendritic spines, from layers II-III of mPFC of rats inoculated with EcoHIV. First, EcoHIV animals exhibited an increased relative frequency of dendritic spines on higher order dendritic branches, which extend superficially into layer I of the mPFC, relative to saline animals; observations which support a profound alteration in synaptic connectivity and/or neurotransmitter innervation. Indeed, monoamine receptor mRNA is widely expressed throughout the mPFC, whereby layers II-III are inundated with serotoninergic, dopaminergic, and adrenergic receptors; only serotonergic mRNA is expressed in layer I (Santana and Artigas, 2017). Second, examination of continuous dendritic spine morphological parameters revealed a profound population shift towards increased dendritic spine head and neck diameter; a morphology consistent with an immature (i.e., stubby: approximately equal dendritic spine head and neck ratio; Peters and Kaiserman-Abramof, 1970) dendritic spine phenotype. Morphological parameters of dendritic spines in pyramidal neurons afford additional evidence for alterations in neurotransmission in the EcoHIV rat; the dendritic spine head receives glutamateric afferents, whereas dopaminergic projections target the dendritic spine neck (Freund et al., 1984). Notably, the inferences drawn from the measurement of dendritic spines are consistent with profound alterations observed in electrophysiological recordings (Cirino et al., 2020), as well as measurements of dopaminergic neurotransmission (for review, McLaurin et al., 2021a; e.g., Kumar et al., 2009; Denton et al., 2019) and glutamatergic homeostasis (for review, Vázquez-Santiago et al., 2014) in HIV-1 seropositive individuals or biological systems utilized to model HIV-1.

The harboring of EcoHIV infection in microglia may partly underlie the profound synaptodendritic alterations observed in the EcoHIV rat. During early HIV-1 infection, the transmigration of infected monocytes and CD4+ cells across the blood-brain barrier leads to the infection of cells, including microglia, in the central nervous system (Cosenza et al., 2002); microglia are also a long-term viral reservoir for productive HIV-1 infection (Ko et al., 2019; Li et al., 2021a). Under homeostatic conditions, microglia are involved in the environmental surveillance of synaptic structures, including dendritic spines (Wake et al., 2009; Tremblay et al., 2010). Microglial aberrations (i.e., activation and/or dysfunction), a consequence of chronic HIV-1 viral protein exposure (Garvey et al., 2014), may lead to and/or result from synaptic dysfunction (for review, Cornell et al., 2022) and alterations in neurotransmission (for review, Domercq et al., 2013; Liu et al., 2016). Furthermore, disruption of microglial specific signaling pathways (i.e., complement receptor 3), which are altered by HIV-1 infection (Mishra et al., 2018), results in persistent synaptic connectivity deficits (Schafer et al., 2012). Indeed, injection of the HIV-1 regulatory protein trans-activator of transcription disrupts microglia-synapse interactions (Tremblay et al., 2013); findings which underscore the critical need to further investigate the relationship between microglial and synaptodendritic dysfunction in HIV-1.

In conclusion, EcoHIV inoculation following a history of cocaine dependence induced profound functional and structural neuroadaptations in the frontal-striatal circuit. From a functional perspective, EcoHIV rats, independent of biological sex, exhibited profound deficits in preference judgment, a simple form of decision-making, and extinction learning. Prominent structural neuroadaptations in frontal-striatal circuitry were evidenced by synaptodendritic alterations in pyramidal neurons from layers II-III of the mPFC. Collectively, the innovative EcoHIV rat affords a fundamental opportunity to system to model frontal-striatal circuit responses to HIV-1 infection following chronic drug use.

## LIMITATIONS OF THE STUDY

Despite the strengths of the present study, a couple of limitations must be acknowledged. First, the pretest-posttest experimental design precluded evaluation of the additive and/or synergistic effect of prior drug use on HIV-1 associated neurocognitive disorders. Second, both drug dependency and EcoHIV inoculation occurred in the mature adult brain. Microglia play a vital role in dendritic (Squarzoni et al., 2014; Lim and Ruthazer, 2021) and synaptic pruning (Paolicelli et al., 2011; Schafer et al., 2012; Mallya et al., 2019), a process that occurs during childhood and adolescence; how the adolescent frontal-striatal circuit responds to HIV-1 infection following drug dependency may be unique. Implementation of different experimental designs would provide an opportunity to address these fundamental questions.

## ACKNOWLEDGEMENTS

This work was supported in part by grants from NIH (National Institute on Drug Abuse, R01- DA013137; National Institute on Drug Abuse, K99-DA056288, National Institute of Mental Health, R01-MH106392; National Institute of Neurological Disorders and Stroke, R01- NS100624). We appreciate the generous gift of the EcoHIV-NL4-3-ECFP lentivirus from Dr. Potash of Icahn School of Medicine at Mount Sinai.

## AUTHOR CONTRIBUTIONS

Conceptualization, C.F.M. and R.M.B.; Methodology, H.L., R.M.B.; Investigation, K.A.M., H.L., S.B.H.; Formal Analysis, K.A.M., C.F.M.; Writing-Original Draft, K.A.M.; Writing-Reviewing & Editing, H.L., C.F.M., S.B.H., R.M.B.; Funding Acquisition, C.F.M., R.M.B.

## STAR METHODS

### RESOURCE AVAILABILITY

#### Lead Contact

Further information and requests for resources should be directed to and will be fulfilled by the lead contact, Rosemarie Booze.

#### Materials Availability

This study did not generate new unique reagents.

#### Data and Code Availability

All relevant data are within the manuscript. Any additional information required to reanalyze the data reported in the manuscript is available from the lead contact upon request.

### EXPERIMENTAL MODEL AND SUBJECT DETAILS

#### Animals

Intact male (*n*=20) and female (*n*=20) Fischer (F344/NHsd; Harlan Laboratories, Inc., Indianapolis, IN) rats were delivered to the animal vivarium between postnatal day (PD) 60 and PD 80. Unrelated animals (i.e., no more than one male and one female from each litter) were requested to preclude violation of the independent observation assumption inherent in many traditional (e.g., analysis of variance (ANOVA)) statistical techniques. Rodent food (Pro-Lab Rat, Mouse, Hamster Chow #3000) and water were available *ad libitum* throughout the duration of the study, unless otherwise specified. Rats were pair- or group-housed with animals of the same sex until jugular vein catheterization (Phase 1.2). Following jugular vein catheterization, animals were single-housed for the remainder of experimentation.

Body weight was evaluated throughout the study to monitor the relative health of EcoHIV and saline animals (Supplementary Figure 3). A statistically significant main effect of week was observed in both male (*F*(22,353)=139.0, *p*≤0.01) and female (*F*(22,353)=112.7, *p*≤0.01) animals; neither a statistically significant Genotype x Week interaction or main effect of Genotype (*p*>0.05) was observed. EcoHIV inoculation, therefore, had no statistically significant effects on somatic growth, consistent with previous observations (Li et al., 2021a).

AAALAC-accredited facilities, with environmental conditions targeted at 21^°^ ± 2^°^C and 50±10% relative humidity, were utilized for the maintenance of animals. The animal colony had a 12-h light: 12-h dark cycle with lights on at 0700h (EST). The project protocol was approved by the Institutional Animal Care and Use Committee (IACUC) at the University of South Carolina (Federal Assurance #D16-00028). Guidelines established by the National Institutes of Health were used for the maintenance of animals and all experimental procedures.

### METHOD DETAILS

#### Apparatus

Sucrose self-administration, cocaine self-administration, and concurrent choice self-administration were evaluated in operant chambers (ENV-008; Med-Associates, St. Albans, VT, USA) located within sound-attenuating enclosures and controlled by Med-PC computer interface software. The front panel of the operant chamber contained a 5 cm x 5 cm opening (ENV 202M- S) through which a recessed 0.01cc dipper cup (ENV-202C) delivered a solution. Additionally, the front panel of the operant chamber had two retractable “active” metal levers (ENV-112BM) and two 28-V white cue lights. The rear panel of the operant chamber had one non-retractable “inactive” metal lever and a 28-V house light. Responding on the “active” lever resulted in reinforcement, whereas responding on the “inactive” lever was not reinforced.

A syringe pump (PHM-100) was utilized to deliver intravenous cocaine infusions through a water-tight swivel (Instech 375/22ps 22GA; Instech Laboratories, Inc., Plymouth Meeting, PA), which was connected to the backmount of the animal using Tygon tubing (ID, 0.020 IN; OD, 0.060 IN) enclosed by a stainless steel tether (Camcaths, Cambridgeshire, Great Britain). Pump infusion times were calculated using a Med-PC computer program based on an animal’s daily bodyweight.

#### Drugs

Cocaine hydrochloride (Sigma-Aldrich Pharmaceuticals, St. Louis, MO), weighed as a salt, was dissolved in physiological saline (0.9%) prior to the start of each testing session to prevent any significant cocaine hydrolysis (Gupta, 1982).

##### Phase 1: Preliminary Training

###### Phase 1.1: Sucrose Self-Administration

*Magazine Training.* Magazine training was conducted for one day to train animals to approach the magazine receptacle from which the sucrose reinforcer was delivered. During the session, a 5% (weight/volume (w/v)) sucrose solution was delivered under a variable interval schedule of reinforcement following the first head poke.

*Autoshaping.* A 42-minute autoshaping procedure was utilized to train animals to lever-press for reinforcement (i.e.. 5% sucrose solution (weight/volume (w/v))) on a fixed-ratio 1 (FR- 1) schedule. The house light was illuminated throughout the duration of the test session. To prevent side bias, an animal was limited to five consecutive presses on a single “active” lever. Successful acquisition of the autoshaping procedure required animals to achieve at least 60 reinforcers for three consecutive or five non-consecutive days.

Thirty-two animals (Male: *n*=15; Female: *n*=17) successfully acquired the autoshaping task within 53 test sessions. Animals (Male: *n*=5; Female: *n*=3) failing to acquire the autoshaping procedure within 63 test sessions were placed on water restriction for 12-15 hours prior to assessment. Two animals (Male: *n*=1; Female: *n*=1) failed to acquire autoshaping within 98 test sessions, despite water restriction, and were not included in subsequent assessments. Water was available *ad libitum* for all subsequent assessments.

*Fixed Ratio Responding: Dose Response.* Following the successful acquisition of autoshaping, male and female animals were assessed using a fixed ratio (FR) schedule of reinforcement, whereby the animal received a reinforcer following each response. After completing a maintenance session (i.e., 5% sucrose on an FR-1 schedule of reinforcement), animals responded for varying sucrose concentrations (i.e., 1, 3, 5, 10, and 30% w/v) during a 42-minute test session. Sucrose concentrations were presented on test days, which occurred every other day, using a Latin-Square experimental design. On intervening non-test days, animals completed a maintenance session.

*Progressive Ratio Responding: Dose Response.* Animals were subsequently assessed using a PR schedule of reinforcement, whereby the response requirement was increased progressively according to the following exponential function: [5e^(reinforcer^ ^number^ ^X^ ^0.2)^]-5 (Richardson and Roberts, 1996). During PR test sessions (Maximum Session Length: 120 Minutes), animals completed the ratio requirement by responding across the two “active” levers. Upon successfully meeting the ratio requirement, the “active” levers were retracted and animals have 4-sec access to one of five sucrose concentrations (i.e., 1, 3, 5, 10, and 30% w/v). Testing days occurred every other day and a maintenance session occurred on the intervening non-test days.

###### Phase 1.2: Jugular Vein Catheterization

Following the completion of sucrose self-administration, a jugular vein catheter was implanted in animals using the methods reported by Bertrand et al. (2018). In brief, 3% inhalant sevofluorane was utilized to induce and maintain anesthesia throughout the surgical procedure. A sterile IV catheter was implanted into the right jugular vein and secured with sterile sutures (4- 0 Perma-Hand silk; EthiconEnd-Surgery, Inc.). The dorsal portion of the catheter/backmount, which connected to an acrylic pedestal embedded with mesh, was implanted subcutaneously above the scapulae and stitched into place (4-0 Monoweb sutures). Post-operative analgesia was provided by subcutaneous administration of butorphenol (Dorolex; 1.0 mg/kg; Merck Animal Health, Millsboro, DE). Infection was prevented via intravenous administration of the antibiotic gentamicin sulfate (0.2 ml 1%; VEDCO, Saint Joseph, MO). Rats were monitored in a heat-regulated warm chamber following surgery and returned to the colony room after recovery from anesthesia.

For seven days after surgery, catheters were flushed daily with a solution containing the anti-coagulant heparin (2.5%; APP Pharmaceuticals, Schamburg, IL) and the antibiotic gentamicin sulfate (1%). Cocaine self-administration began five days after jugular vein catheterization. Prior to operant testing each day, catheters were flushed with 0.9% saline solution (Baxter, Deerfield, IL). After the completion of operant testing, catheters were flushed with post-flush solution containing heparin (2.5%) and gentamicine sulfate (1%).

###### Phase 1.3: Cocaine Self-Administration

*Fixed Ratio Responding.* First, during a 60-minute test session, rats responded for a 0.33 mg/kg/infusion of cocaine according to a FR-1 schedule of reinforcement. FR-1 assessments were conducted for five days.

*Progressive Ratio Responding.* Second, rats responded for a 0.75 mg/kg/infusion of cocaine on a PR schedule of reinforcement, whereby the response requirement was defined using the exponential function presented above (Phase 1.1). PR assessments, which lasted a maximum of 120-minutes, were conducted for seven days.

##### Phase 2: Pretest Evaluation

###### Phase 2.1: Concurrent Choice Self-Administration

After establishing a history of both sucrose and cocaine self-administration, male and female animals were evaluated in a concurrent choice self-administration experimental paradigm for five days. During the 60-minute assessment of choice behavior, animals were responding on an FR-1 schedule of reinforcement for either 5% sucrose (w/v) or 0.33 mg/kg/infusion of cocaine. The position (i.e., right or left) of the sucrose-paired lever was randomly assigned.

At the beginning of each assessment, animals completed four forced choice trials, (i.e., only one “active” lever was available) for two sucrose and two cocaine reinforcers. Subsequently, both “active” levers were simultaneously available to allow the animals to freely choose between sucrose and cocaine. A 20-second time-out occurred after each “active” lever response.

##### Phase 3: EcoHIV (i.e., Posttest) Evaluation

###### Phase 3.1: Bilateral Retro-Orbital EcoHIV or Saline Injections

*EcoHIV Virus Construction and Preparation.* The EcoHIV-NL4-3-EGFP lentivirus, kindly provided by M.J. Potash (Icahn School of Medicine at Mount Sinai), was constructed from pNL4-3 and pNCA-WT. The fragment of NL4-3 at nucleotide 6310, NL4-3 at nucleotide 7751, NCA-WT at nucleotide 6229, and NCA-WT at nucleotide 8223, and were ligated to generate the chimeric virus. Plasmid DNA was transfected into 293FT cells (Lipofectamine^TM^ 3000, Cat. No. L3000015, Invitrogen) and virus stocks were prepared from the conditional medium. Virus stocks were subsequently concentrated using the Lenti-X concentrator (Cat. No. 631231, Clontech Laboratories, Mountain View, CA, USA).

*Treatment Assignment.* Animals were assigned to receive either EcoHIV or saline injections based on three measurements from preliminary training, including: 1) the number of days to acquire autoshaping; 2) the average number of “active” lever presses across the seven days of cocaine PR responding; and 3) concurrent choice self-administration preference (i.e., No Choice, Sucrose or Cocaine). Treatment assignment yielded sample sizes of EcoHIV, *n*=20 (male, *n*=10, female, *n*=10) and Saline, *n=18* (male, *n*=9, female, *n*=9).

*Bilateral Retro-Orbital Injections.* Male and female rats were anesthetized with inhalant sevofluorane and positioned laterally with the injection-eye facing up. A 1cc tuberculin syringe with a 26G needle was slowly inserted into the medial canthus of the eye into the retro-orbital vessels. 20 μL of either EcoHIV NL4-3-EGFP (2.809 x 106 TU/mL) or saline was gently injected into the retro-orbital vessels. The procedure was repeated in the other eye. After bilateral inoculation, the rat recovered in a heat-regulated warm chamber.

*Immunohistochemical Staining.* After the completion of experimentation, immunohistochemical staining was utilized for two complementary purposes, including 1) to quantity the amount EcoHIV infection present in the mPFC; and 2) to establish the predominant cell type expressing EcoHIV infection. Following transcardial perfusion and removal of the animals’ brain, tissue was post-fixed in cold paraformaldehyde (4%). A vibratome (PELCO easiSlicer^TM^, TED PELLA, INC., Redding, CA, USA) was utilized to section brain tissue into 50 μm thick coronal slices. Brain tissue sections were incubated overnight (4°C) with mouse monoclonal anti-GFP antibody (Cat #: ab1218, abcam, Boston, MA) and Alexa Fluor 697 rabbit monoclonal anti-Iba1 antibody (Cat #: ab225261, abcam) to stain for EcoHIV-EGFP and microglia, respectively. A Nikon TE-2000E confocal microscope system was used to obtain z-stack images (Z-plane interval: 0.15 μm) with a 60x oil objective (*n.a.*=1.4). EcoHIV expression was quantified by counting the number of EGFP signals within a representative area.

###### Phase 3.2: Concurrent Choice Self-Administration (3 to 9 Days Post Inoculation)

Beginning three to four days after bilateral retro-orbital injections of either EcoHIV or saline, animals were reassessed (i.e., first posttest evaluation) in the concurrent choice self-administration paradigm (described in Phase 2.1: Concurrent Choice Self-Administration) for five days.

###### Phase 3.3: Sucrose Self-Administration (8 to 22 Days Post Inoculation)

The reinforcing efficacy of sucrose in EcoHIV and saline animals was subsequently assessed using the PR dose-response experimental paradigm described in Phase 1.1.

###### Phase 3.4: Cocaine Self-Administration (21 to 39 Days Post Inoculation)

A PR dose-response experimental paradigm was utilized to evaluate the reinforcing efficacy of cocaine in EcoHIV and saline animals. Rats responded according to a PR schedule of reinforcement, whereby the response requirement was defined using the exponential function presented in Phase 1.1, for one of six cocaine concentrations (i.e., 0.01, 0.03, 0.10, 0.33, 0.75, and 1.0 mg/kg/infusion). Cocaine concentrations were presented in an ascending manner.

Testing days occurred every other day and a maintenance session (i.e., 0.33 mg/kg/infusion of cocaine on an FR-1 schedule of reinforcement) occurred on the intervening non-test days. One saline female animal died during the cocaine self-administration task.

###### Phase 3.5: Concurrent Choice Self Administration (37 to 45 Days Post Inoculation)

Beginning 37 to 40 days after bilateral retro-orbital injections of either EcoHIV or saline, animals were reassessed (i.e., second posttest evaluation) in the concurrent choice self-administration experimental paradigm (described in Phase 2.1: Concurrent Choice Self-Administration) for five days.

###### Phase 3.6: Extinction (42 to 52 Days Post Inoculation)

Following completion of the second posttest evaluation, each animals’ preferred reinforcer (i.e., Cocaine or Sucrose) was individually determined and replaced (i.e., Cocaine with Saline; Sucrose with Water) to evaluate extinction learning in the concurrent choice self-administration experimental paradigm (described in Phase 2.1: Concurrent Choice Self-Administration). Cocaine extinction learning yielded sample sizes of EcoHIV, *n*=14 (male, *n*=7, female, *n*=7) and Saline, *n=*10 (male, *n*=7, female, *n*=3). Saline extinction learning yielded the following sample sizes: EcoHIV, *n*=6 (male, *n*=3, female, *n*=3) and Saline, *n=*6 (male, *n*=1, female, *n*=5). One saline male animal failed to respond during the second posttest evaluation and did not complete extinction learning. Extinction learning was evaluated for seven days.

##### Phase 4: Neuroanatomical Assessments

###### Phase 4.1: Synaptodendritic Alterations in Pyramidal Neurons in the Medial Prefrontal Cortex

*Estrous Cycle Tracking.* Estrous cycle stage was determined immediately prior to sacrifice using cellular cytology in vaginal smears under a 10x light microscope. To decrease potential hormonal variability, the goal was to sacrifice female EcoHIV and saline animals during the diestrus phase.

*Tissue Preparation.* EcoHIV and saline animals were deeply anesthetized (5% sevoflurane; Abbot Laboratories, North Chicago, IL, USA) and transcardially perfused. After removal, the rat brain was placed in 4% paraformaldehyde for 10 min before being sliced coronally (500 μm) using a rat brain matrix (ASI Instruments, Warren, MI, USA). Coronal brain slices were placed into a tissue cell culture plate (24 well-plate; Corning, Tewksbury, MA, USA) with 1mM phosphate-buffered saline (PBS). PBS was removed from the targeted wells before ballistic labeling.

*Ballistic Labeling.* The ballistic labeling protocol published by Li et al. (2020) was utilized to evaluate synaptodendritic alterations in pyramidal neurons in the mPFC. Tefzel tubing was filled with a polyvinylpyrrolidone (PVP) solution, which included 100 mg of PVP dissolved in 10 mL ddH_2_O, for 20 minutes; the PVP solution was subsequently expelled from the Tefzel tubing. Ballistic cartridges were prepared by thoroughly mixing 170 mg of tungsten beads (Bio-Rad, Hercules, CA, USA) in 250 μL of 99.5% methylene chloride (Sigma-Aldrich, St. Louis, MO, USA). Additionally 6 mg of DiIC19(3) dye (Invitrogen, Carlsbad, CA, USA) were dissolved in 300 μL of 99.5% methylene chloride. The tungsten bead suspension was pipetted onto a glass slide and allowed to air dry. The DiIC19(3) dye mixture was subsequently added on top of the tungsten bead suspension layer; the two layers were mixed thoroughly and split into two centrifuge tubes (1.5 mL) filled with ddH_2_O. The tungsten bead/DiIC19(3) dye mixture was homogenized using sonication, slowly drawn into the PVP coated Tefzel tubing, and placed into the tubing preparation station (Bio-Rad). After rotating for one minute, all water was removed from the PVP coated Tefzel tubing; rotation was subsequently continued for 30 minutes under nitrogen gas (0.5 L per minute). Ballistic cartridges were cut into 13 mm lengths and stored in the dark.

The Helios gene gun (Bio-Rad), which had a piece of filter paper between the two mesh screens, was loaded with the prepared ballistic cartridges and connected to the helium hose (Output Pressure: 90 pounds per square inch). The applicator was placed vertically at the center of the targeted tissue well and approximately 1.5 cm away from the brain tissue. Tissue was washed three times with 100 mM PBS and stored at 4°C in the dark for three hours before being mounted on a glass slide using Pro-Long Gold Antifade (Invitrogen).

*Confocal Imaging.* A confocal microscopy system (Nikon TE-2000E and Nikon’s EZ-C1 software (version 3.81b)) was utilized to obtain z-stack images (60x Oil Objective, *n.a.*=1.4, Z-Place Interval of 0.15 μm) of pyramidal neurons in the mPFC, located approximately 3.7 mm to 2.2 mm anterior to Bregma (Paxinos and Watson, 2014). Three neurons per animal were imaged.

*Neuronal and Dendritic Spine Analysis.* Neurolucida 360 (MicroBrightfield, Williston, VT, USA), a sophisticated neuronal reconstruction software, was utilized to analyze pyramidal neurons, and associated dendritic spines, from the mPFC. After blinding, one neuron was selected from each animals based on stringent selection criteria (e.g., continuous dendritic staining, low background/dye clusters; Li et al., 2020), yielding the following sample sizes: EcoHIV, *n*=20 (male, *n*=10, female, *n*=10) and Saline, *n=*17 (male, *n*=9, female, *n*=8).

Pyramidal neuron morphology was evaluated using a centrifugal branch ordering scheme, a measure of dendritic branching complexity. Dendritic spine morphology was characterized by three parameters, including dendritic spine backbone length (μm), head diameter (μm), and neck diameter. Based on previously published results, boundary conditions for dendritic spines were established (volume, 0.05 to 0.85 µm: Radley et al., 2008; backbone length, 0.4 to 4.0 µm: Arellano et al., 2007, Ruszczycki et al., 2012; head diameter, 0 to 1.2 µm: Konur et al., 2003); dendritic spines failing to meet any of the boundary conditions were excluded from the analysis.

## QUANTIFICATION AND STATISTCAL ANALYSIS

Statistical approaches, including analysis of variance (ANOVA) and regression techniques, were utilized to analyze all behavioral and neuroanatomical data (SAS/STAT Software 9.4, SAS Institute Inc., Cary, NC, USA; SPSS Statistics 27, IBM Corp., Somer, NY, USA; GraphPad Prism 5 Software Inc., La Jolla, CA, USA). Figures were created using GraphPad Prism 5 Software Inc. (La Jolla, CA, USA). To preclude any potential compound symmetry violations, orthogonal decompositions or the conservative Greenhouse-Geisser *df* correction factor (*p*_GG_; Greenhouse and Geisser, 1959) were implemented. Complex interactions were futher evaluated using simple effects. An alpha level of *p*≤0.05 was set for the establishment of statistical significance.

Repeated measures ANOVA techniques (PROC MIXED; SAS/STAT Software 9.4) were utilized to statistically analyze body weight, PR responding during cocaine escalation, sucrose dose response, and cocaine dose response, as well as data from the concurrent choice self-administration task. Between-subjects factors included genotype (EcoHIV vs. Saline) and biological sex (Male vs. Female), whereas within-subjects factors included time (e.g., day, week), concentration, dose, or reinforcer (i.e., Sucrose vs. Cocaine) as appropriate. A compound symmetry covariance structure was utilized for analyses conducted in PROC MIXED. For the concurrent choice self-administration task, a choice was defined as responding for at least five reinforcers total; if an animal failed to respond for at least five total reinforcers, the data were censored in both the figure and analysis.

Extinction learning was also analyzed using a repeated measures ANOVA (SPSS Statistics 27). Day, which included baseline (i.e., Day 5 of the Second Posttest Evaluation of Preference Judgment) and the seven days of extinction, served as the within-subjects factor. Genotype (EcoHIV vs. Saline) and the reinforcer removed (i.e., Sucrose vs. Cocaine) served as between-subjects factors. Due to sample size limitations, biological sex was not included as a between-subjects factor.

A generalized linear mixed-effects model, conducted using PROC GLIMMIX (SAS/STAT Software 9.4), with a random intercept and Poisson distribution was utilized to statistically analyze all neuroanatomical data. An unstructured covariance structure was utilized for analyses conducted in PROC GLIMMIX. Genotype (EcoHIV vs. Saline) and biological sex (Male vs. Female) were included as between-subjects factors. Branch Order or Bin were served as within-subjects factors, as appropriate.

