## Supplementary Information for "Disrupted Decision-Making: EcoHIV Inoculation Following a History of Cocaine Use"

Cognitive and Neural Science Program

Department of Psychology

University of South Carolina

Columbia SC 29208

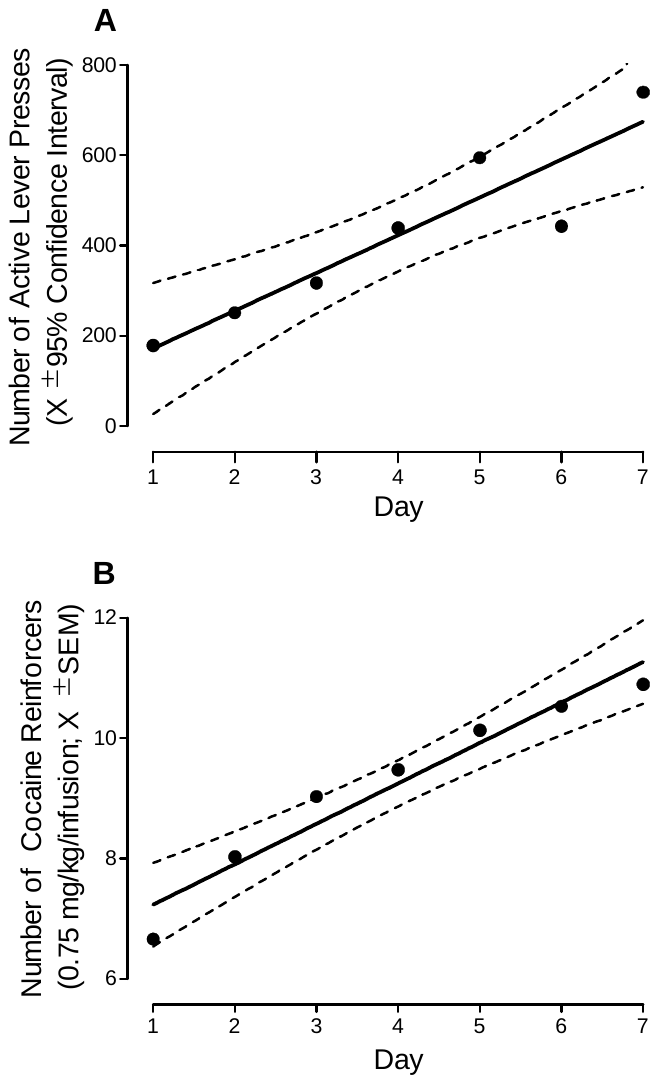

**Figure S1: Cocaine Escalation.** Across seven progressive ratio test sessions, animals exhibited a linear increase in the number of active lever presses (**A**) and the number of cocaine reinforcers (**B**) supporting the development of a drug dependent phenotype.

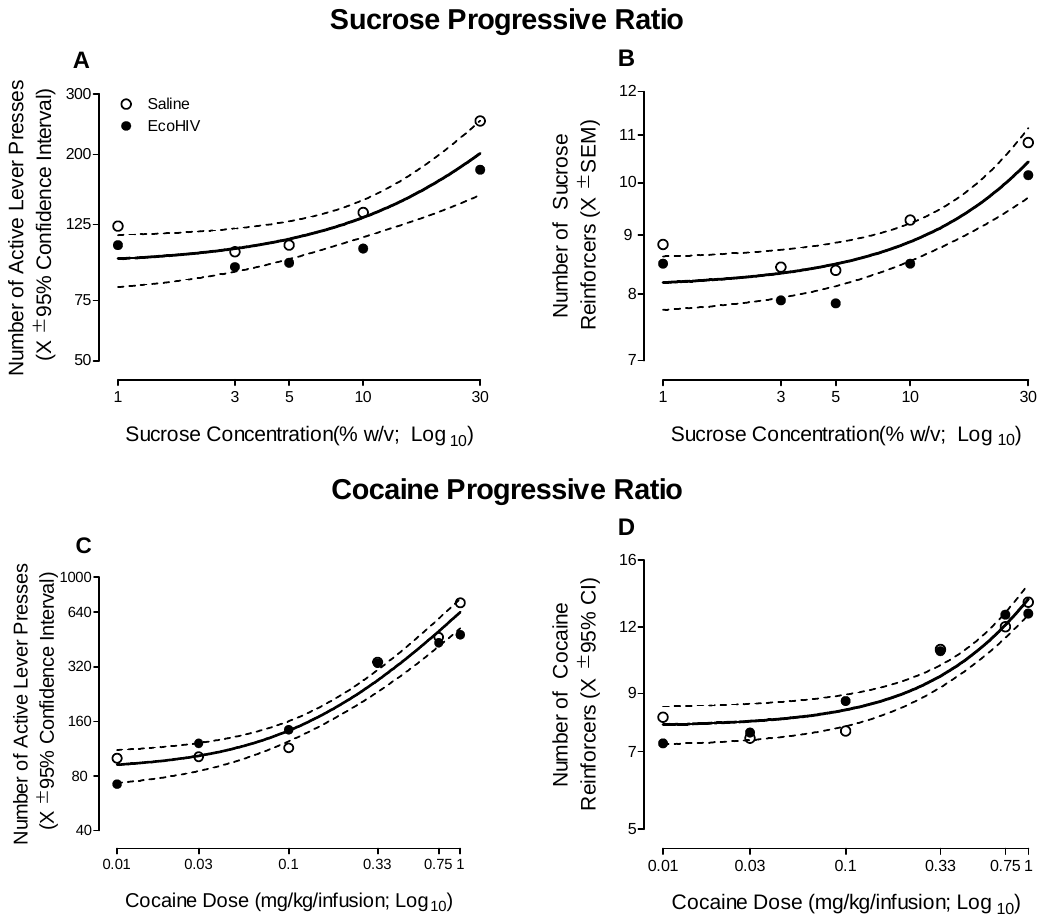

**Figure S2: Reinforcing Efficacy of Sucrose and Cocaine.** A progressive ratio dose-response experimental paradigm was used to evaluate the reinforcing efficacy of sucrose and cocaine in EcoHIV and saline animals. EcoHIV inoculation failed to alter the reinforcing efficacy of either sucrose (**A-B**) or cocaine (**C-D**) evidenced by global best-fit functions for both the number of active lever presses and number of reinforcers.

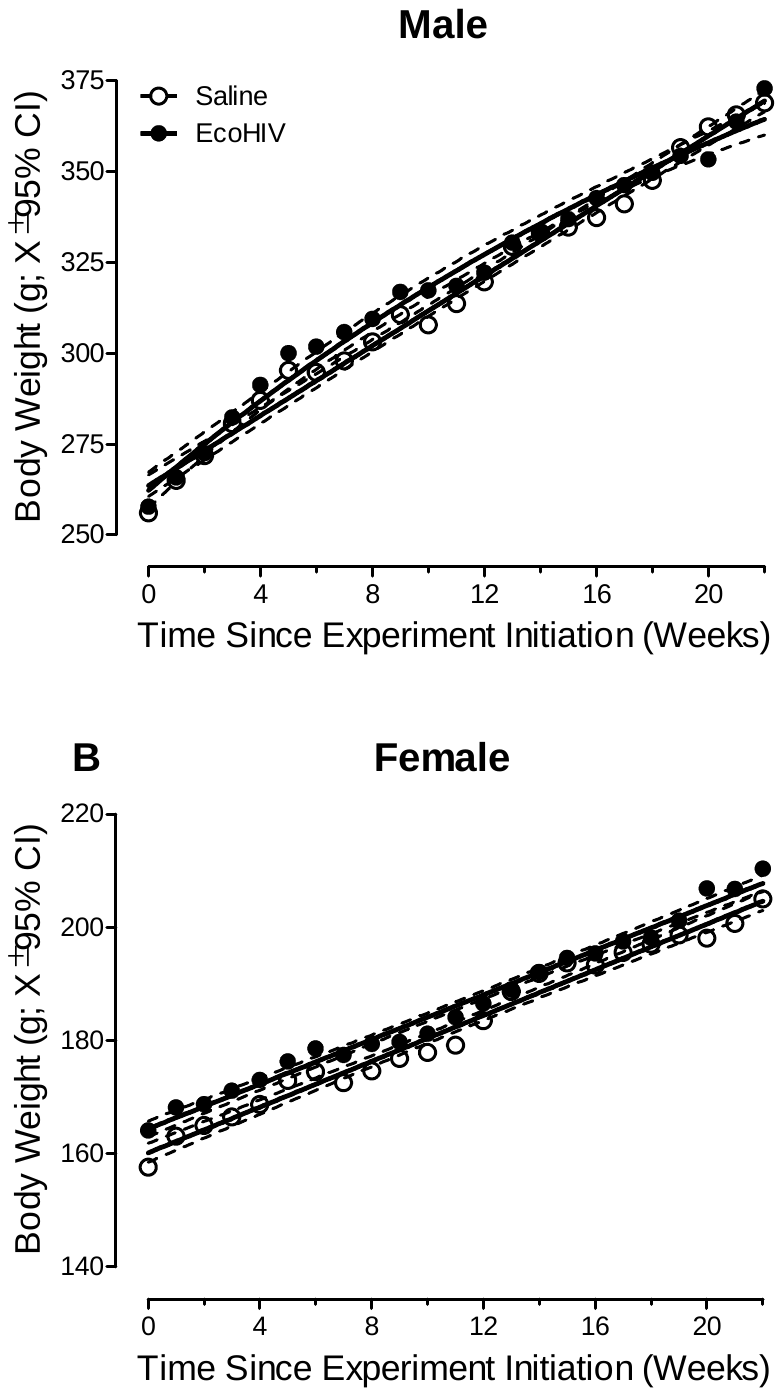

**Figure S3: Body Weight.** Mean body weight, a measure of somatic growth, is illustrated for male (**A**) and female (**B**) animals as a function of genotype (EcoHIV vs. Saline). All animals, independent of genotype, exhibited significant growth throughout the duration of the study. EcoHIV inoculation, therefore, had no adverse effects on somatic growth.
